# The fidelity of mRNA translation as a novel regulatory layer for brain development

**DOI:** 10.64898/2026.04.01.715671

**Authors:** Jacob A Best, Pitchaporn Akaphan, Indu Tripathi, Ashakiran B Rochette, Riku Nagai, Kryshna C Miller, Andrea Suárez, Lien Nguyen, Kotaro Fujii

## Abstract

Although the fidelity of mRNA translation is essential for maintaining proteome integrity, whether translation error rates vary across cell types and developmental stages in vivo remains largely unexplored. Here, we generate a gain-of-activity dual-luciferase knock-in reporter mouse that enables quantitative monitoring of translation errors in vivo. Using this system, we systematically characterize the spatiotemporal dynamics of translation fidelity across mammalian development. Mature organs exhibit lower error rates than pluripotent embryonic stem cells. Translation fidelity diverges sharply among organs, becoming progressively established during embryonic development, with brain and muscle displaying the highest accuracy. To determine functional significance, we experimentally increased translation errors during cerebral organoid formation and in vitro neuronal differentiation. Elevated error rates reduced neuronal output by impairing neuronal maturation without altering neural progenitor populations. Consistently, differentiated neurons display uniformly elevated fidelity across multiple classes of translation errors, including stop codon readthrough, amino acid misincorporation, and ribosomal frameshifting. These findings demonstrate that translation fidelity is not a fixed intrinsic property of the translation machinery but is developmentally regulated and required for efficient neuronal differentiation. Together, our results identify translation fidelity as a developmentally tuned, tissue-specific dimension of gene expression in vivo.

## Introduction

Establishing tissue and cell-type-specific proteomes at the correct time and place is essential for coordinating cell differentiation and migration during embryonic development. While quantitative regulation of gene expression has been extensively studied, how the quality of gene expression, including the fidelity of mRNA translation, changes between cell types and contributes to developmental processes remains poorly understood. Despite the importance of maintaining mRNA translation fidelity ^1,2^, mRNA translation is inherently error-prone with errors occurring at an estimated rate of ∼10⁻³–10⁻⁴ per codon ^3^. Approximately 15% of average-length proteins contain at least one decoding error ^4^. It remains largely unexplored whether this high level of translation error simply reflects a fixed biochemical limitation of the translation machinery, manifesting as stochastic noise, or instead represents a dynamically regulated layer of gene expression across tissues and developmental stages.

Although in vitro biochemical and single molecule studies have provided detailed insights into the kinetics of ribosomal misdecoding ^5–8^, quantifying translation errors in vivo remains challenging. Ribosome profiling (Ribo-seq) has revolutionized our understanding of translation regulation, including in the context of embryonic development ^9–13^, yet its ability to capture translation errors is limited. Mass spectrometry (MS) can, in principle, identify error-containing peptides, but such products are rare and difficult to distinguish from analytical misassignments, often necessitating stringent validation and integration across datasets ^14,15^. In contrast, gain-of-function dual-luciferase reporters, which contain an active-site mutation within the luciferase gene, can be used to detect amino acid misincorporation with high sensitivity and low background readout ^16,17^. Similarly, a stop codon or frameshift element can be inserted between tandem luciferase coding sequences to monitor stop codon readthrough or ribosomal frameshifting ^18,19^. Both MS and gain-of-function reporter approaches indicate that most amino acid misincorporation and stop codon readthrough events occur with near-cognate errors, suggesting these errors occur during ribosomal decoding rather than from tRNA mis-aminoacylation ^15,20,21^.

Translation error-containing proteins are prone to misfolding and can be eliminated by the ubiquitin-proteasome system ^22^. Indeed, nearly one-third of newly synthesized proteins exhibit rapid degradation kinetics within 10 minutes ^23–25^. Thus, protein quality control mechanisms maintain proteome integrity despite the continuous production of error-containing proteins. However, increased translation errors can destabilize this balance and compromise proteostasis. Indeed, increasing mis-aminoacylation has been shown to increase the number of unfolded proteins and induce neurodegeneration ^26^. Furthermore, mice with error-prone ribosomes are known to exhibit accelerated aging phenotypes, especially in the skeletal muscle and brain ^27–29^. Consistent with the link between translation accuracy and tissue homeostasis, translation fidelity has been associated with lifespan in yeast, fly, worm, and rodent ^30–33^. Notably, increases in stop codon readthrough during aging in skeletal muscle and brain have been demonstrated using a gain-of-function fluorescent-luciferase reporter knock-in mouse model ^34^.

Given the intimate links between translation fidelity, tissue vulnerability, and organismal lifespan, the mechanisms that establish fidelity during embryonic development may have long-lasting consequences for organismal health. However, whether translation fidelity is dynamically programmed across tissues during embryogenesis remains unknown. Although translation errors are generally viewed as detrimental due to their impact on proteome stability, emerging evidence suggests that error-derived peptides can have beneficial effects in specific contexts ^35–39^. This raises the possibility that context-dependent modulation of translation fidelity may influence proteome diversity or cellular adaptability under physiological and pathological conditions.

Here, using a highly quantitative reporter knock-in mouse model, we uncover unexpected spatiotemporal dynamics of translation fidelity during organ development. We find that stop codon readthrough occurs at substantially lower levels in intact organisms than in pluripotent stem cells and decreases progressively across development, revealing a developmental tightening of translational accuracy as organs mature. Notably, the brain and muscle tissues exhibited the lowest error rates. Particularly, increasing translation errors impaired neuronal differentiation in both cerebral organoid and neuron differentiation models. Together, we uncovered that mRNA translation fidelity is a developmentally programmed layer of gene regulation.

## Results

### Development of a highly quantitative reporter knock-in mouse

To delineate the spatiotemporal dynamics of translation fidelity in organisms, we generated a knock-in mouse model that harbors a dual-luciferase gain-of-activity reporter in its genome. While dual-luciferase reporters possess a wide dynamic range and high sensitivity, it is difficult to monitor both Renilla (Rluc) and Firefly (Fluc) luciferases under the microscope or by cell sorting. Thus, we added both peptide and enzymatic tags to the dual luciferase. Our reporter system contains Flag-Rluc-SNAP and Halo-Fluc-HA cassettes in tandem with the UGA stop codon between them. Accurate mRNA translation produces only Flag-Rluc-SNAP, while both Flag-Rluc-SNAP and Halo-Fluc-HA are produced when the ribosome reads through the stop codon. To quantify stop codon readthrough, the ratio of Fluc to Rluc activity was monitored using Rluc as the normalizer to count errors per mRNA translation event (Fig. 1A). To generate a knock-in mouse model, we started with UGA stop codon readthrough, which has the highest stop codon readthrough rate in cultured cells ^40^. Transfection of our reporter into cultured mouse embryonic stem cells (mESCs) was able to recapitulate this result (Fig. 1B). Different readthrough rates between stop codons, together with the substantially lower transcriptional error frequency (∼10⁻⁶ in yeast) ^41^, indicate that the observed signal is most consistent with ribosomal misdecoding. To visualize our reporter, we employed the Halo and SNAP enzymatic tags, which are able to form covalent bonds with specific fluorophore-ligated ligands ^42,43^. Halo-tag treated with a synthetic ligand conjugated with the Janelia Fluor 646 ^44^ was able to visualize small amounts of error products at single-cell resolution with transient transfection of our reporter constructs in mESCs (Figs. 1C, and S1). The control SNAP tag was also visualized by Oregon green ligand ^45^, indicating a heterogeneous amount of Halo-tag containing products between individual cells within the same mESC population (Fig. 1C). However, the visualization of error products using Halo tag ligands has not succeeded with knock-in mouse tissue sections or derived cells due to the small number of error-containing products.

**Fig. 1:**
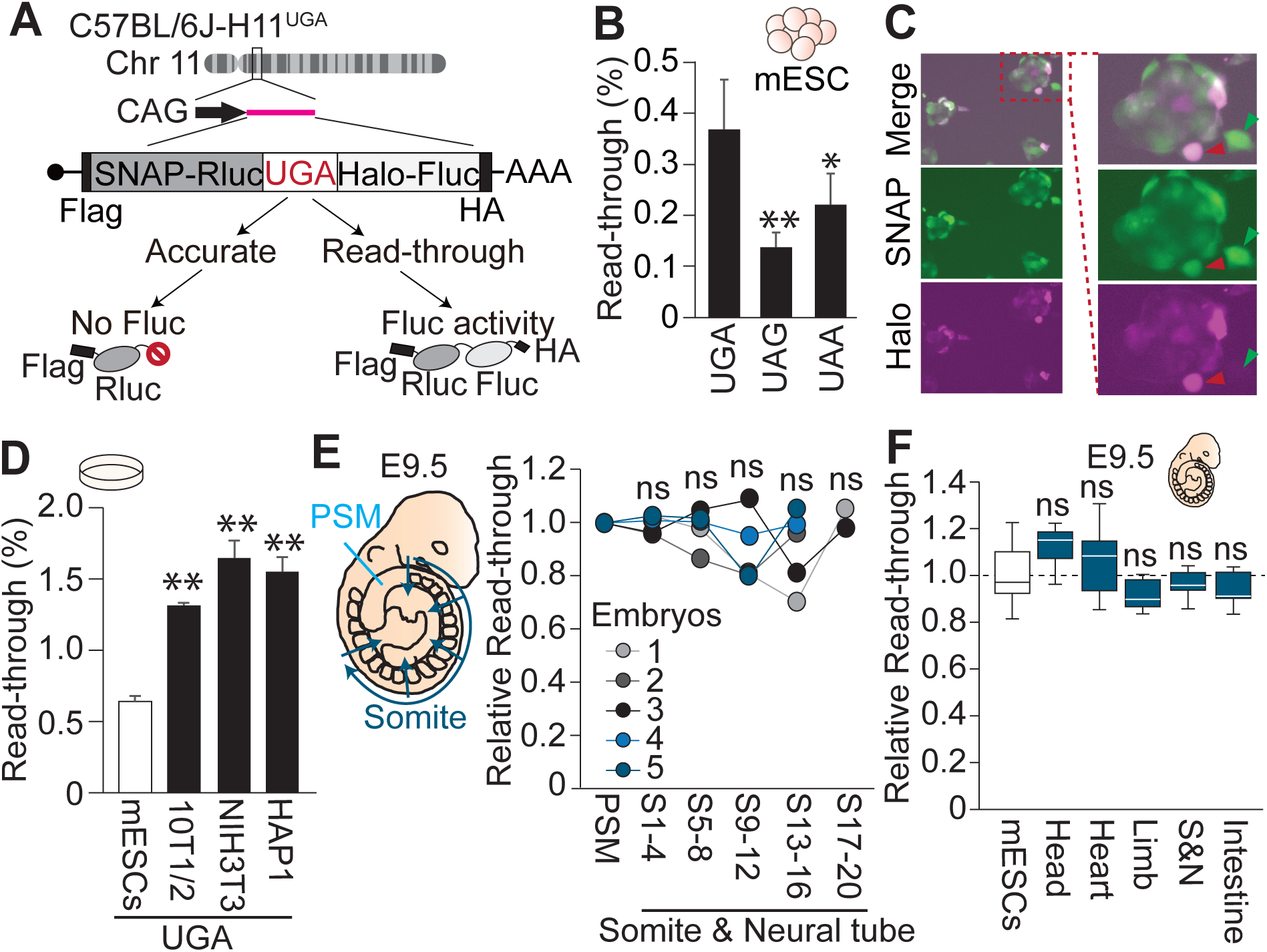
Novel gain-of-activity reporter system monitoring stop codon readthrough indicates no spatial differences in stop codon readthrough errors in embryonic day 9.5. **(A)** Schematic of the knock-in mouse model monitoring UGA stop codon readthrough in vivo. Dual-luciferase gain-of-activity reporter was inserted into the *H11* locus of the C57BL/6J mouse genome. Reporter mRNA was transcribed from the CAG promoter. When the ribosome reads through the UGA stop codon, the Halo-Fluc domain will be translated to quantitatively compare stop codon readthrough products between samples. **(B)** Each stop codon exhibited significantly different readthrough rates compared to the UGA stop codon with our reporter system by transient transfection into mouse embryonic stem cells (mESCs). Read-through rates were calculated based on the value of non-stop construct. Data are graphed as mean ± SD (N>3). *P<0.05, **P<0.01 by unpaired t test. **(C)** Heterogeneity of readthrough products was observed between cells in transiently transfected mESCs by visualizing Halo and SNAP tags. The green triangle indicates a cell with less error product, and the red triangle indicates a cell with more error product. **(D)** Immortalized cell lines exhibited higher UGA readthrough rates than mESCs. Data are graphed as mean ± SD (N>3). **P<0.01 by unpaired t test. **(E)** Error rate did not change across somite and neural tube differentiation in E9.5. Presomatic mesoderm (PSM) and each four somite and neural tube segments are microdissected and monitor readthrough products for each embryo. Embryos were harvested from 2 litters. No segments indicated significant differences by paired t test. **(F)** All tissues at E9.5 exhibited comparable UGA stop codon readthrough products with mESCs. Box plots show the interquartile range (IQR) with the median indicated by a central line; whiskers extend to 1.5 × IQR. n≥10 harvested from more than 2 litters. ns: not significant by Mann-Whitney/Kruskal-Wallis.

To achieve systemic and high-level expression, we integrated the reporter transgene with the CAG promoter into the *Hipp11* locus of the C57BL/6J mouse ^46,47^. To evaluate if the transgene resulted in any phenotypic abnormalities, we monitored mouse growth by recording weight from the day of birth, postnatal day 0 (P0), to 22 weeks old. Knock-in and WT mice exhibited similar growth rates in both male and female without any notable maturation defects or abnormalities (Fig. S2A). To validate the ability of our knock-in reporter to detect changes in mRNA translation errors, we generated mouse embryonic fibroblast (MEF) lines from our knock-in mice and treated the cells with paromomycin, an aminoglycoside antibiotic, which induces stop codon readthrough and AA misincorporation ^48–50^. Paromomycin directly binds the 40S decoding center at the A-site and stabilizes mis-matched codon-anticodon pairings with near-cognate tRNA. As expected, MEF lines with paromomycin-treated cells exhibited significantly more Fluc activity than non-treated cells without concurrent changes in Rluc activity (Fig. S2B). Thus, our integrated reporter was able to quantify the expected changes in stop codon readthrough error rate under paromomycin treatment, indicating that our knock-in mice are suitable to monitor spatiotemporal changes in translation fidelity.

### Each organ has a different stop codon readthrough rate in young adult mice

Given that mESCs have lower translation errors than other immortalized cell lines (Fig. 1D), we initially thought that pluripotent stem cells and early-stage embryos would have the highest fidelity compared to mature tissues. Hence, we hypothesized that cells and tissues establish specialized fidelity across differentiation, either maintaining fidelity or increasing translation errors. To examine our hypothesis, we began by monitoring translation error across the differentiation process from presomitic mesoderm (PSM) to posterior somites, which are produced during axis elongation, and to anterior somites, which undergo differentiation into sclerotome and dermomyotome in response to developmental signaling cues ^51^. In parallel with somite maturation, the neural tube also displays a posterior-anterior differentiation gradient, where anterior regions are fully patterned dorsoventrally while posterior regions near the PSM remain immature. Previous work has shown significant mRNA translation regulation during mesoderm and neural tube differentiation, particularly within the developmental signaling pathway components ^11,52^, while the impact and dynamics of translation fidelity have not been explored. To examine if mRNA translation fidelity changes dynamically across cell differentiation, we microdissected the PSM and the neural tube together with the paired somites at each axial level, collecting tissues in four-somite blocks from posterior to anterior. To our surprise, we did not observe differences in stop codon readthrough rate across the anterior-posterior axis of the somite and neural tube (Fig. 1E). Furthermore, we also examined stop codon readthrough across different body regions at embryonic day 9.5 (E9.5), including the head, somite-neural tube, heart, limb bud, and developing intestine. Strikingly, on E9.5, all tissues possessed a stop codon readthrough rate similar to that of mESCs (Figs. 1F, and S2C), indicating that stop codon readthrough rate is largely homogenous at this stage of the developing embryo.

To further explore if translation fidelity changes between organs, we harvested organs from 2-month-old young adult mice after transcardiac perfusion to minimize blood contamination. We harvested the brain, heart, quadriceps (skeletal muscle), lung, liver, and kidney. In contrast to E9.5, all the organs we tested indicated lower stop codon readthrough rate than mESCs in both females and males (Figs. 2A, and S2D). Contrary to our hypothesis that mESCs exhibit higher translation fidelity than mature tissues, these results suggest that in vivo organisms display higher translation fidelity than cultured cells. To test this hypothesis, the stop codon readthrough rate was monitored across primary MEF line derivation. We observed that translation errors quickly increased within a single passage (Fig. 2B), suggesting there could exist a pressure on cells to maintain higher fidelity in vivo that is relaxed when their environment shifts to in vitro conditions.

**Fig. 2:**
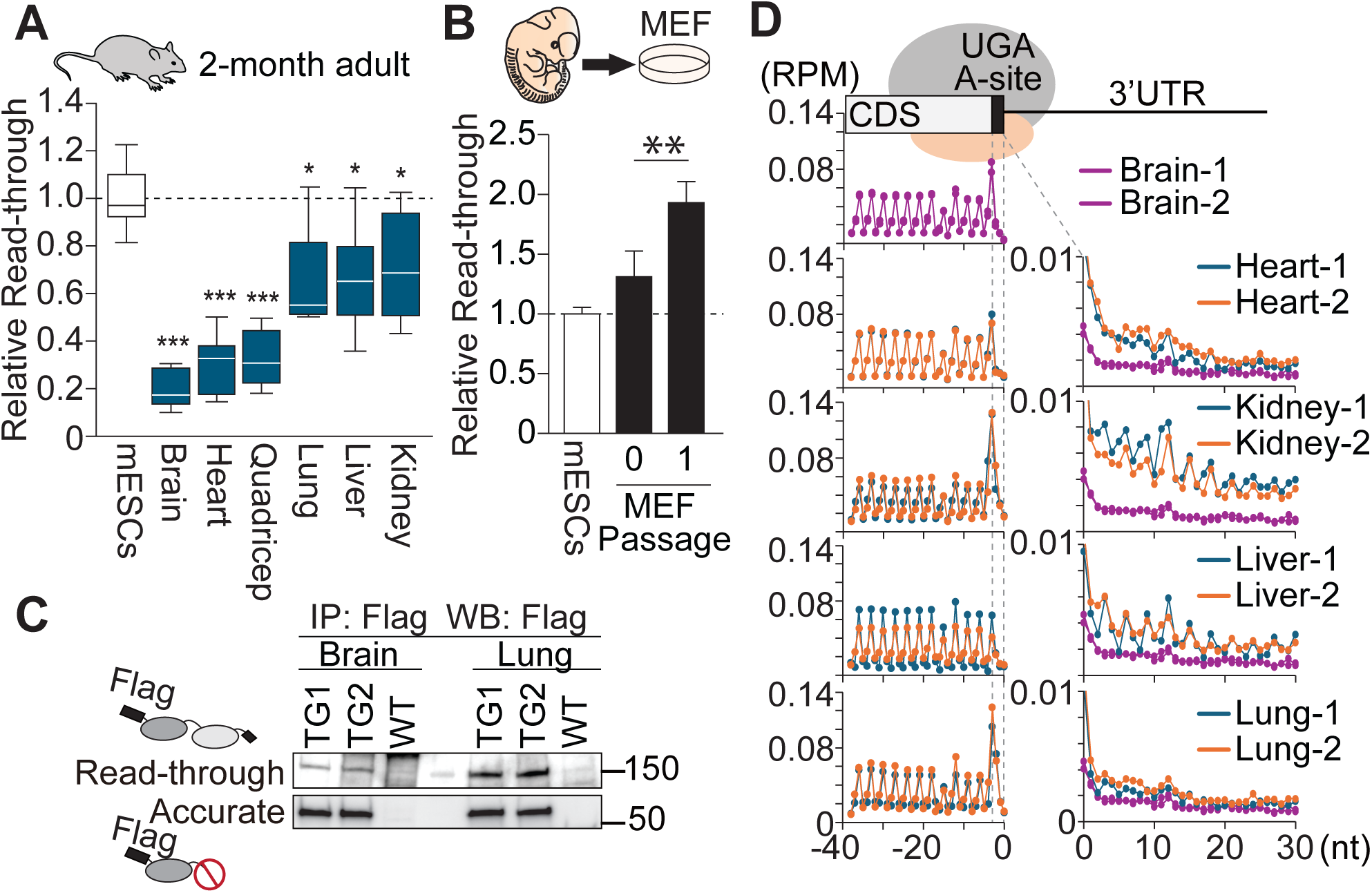
Mature organs exhibit distinct translation error rates. (**A**) Knock-in mouse model indicated different stop codon readthrough rates among organs in 2-month-old mice. The relative stop codon readthrough rate in each organ compared to mouse embryonic stem cells (mESCs) was quantified from ≥3 males and ≥3 females. Box plots show the interquartile range (IQR) with the median indicated by a central line; whiskers extend to 1.5 × IQR. *P<0.05, **P<0.01, ***P<0.001 by Mann-Whitney U test. (**B**) Stop codon readthrough products increased within a single passage of mouse embryonic fibroblast (MEF). n≥3 **P<0.01 with unpaired t test. (**C**) Western blot against reporter proteins indicated higher readthrough products in the lung than in the brain. Reporter proteins were immunoprecipitated (IP) using anti-Flag antibodies, and then western blotting (WB) was performed using anti-Flag. Accurately translated products (Accurate) are distinguished and separately detected from stop codon readthrough products (Read-through). (**D**) Ribosome profiling (Ribo-seq) shows a genome-wide trend of less UGA stop codon readthrough with endogenous transcripts in the brain. Datasets from a previously published 7-week-old young adult mouse were used. Genome-wide analysis of A-site position indicates a triplet pattern in the coding sequence (CDS) on the left. The right side of the plot shows ribosome density immediately after the UGA stop codon in the 3’ untranslated region (3’ UTR). Y-axis plotted in reads per million (RPM). The UGA stop codon was aligned from −3 to −1.

Our results indicate that the brain, heart, and quadriceps exhibited the lowest stop codon readthrough rate among the organs harvested (Fig. 2A). To validate our results shown in the luciferase assay, we also conducted western blotting to compare error products to accurate products between tissues. Error and accurate products were immunoprecipitated using an antibody against N-terminal Flag peptides from the brain and lungs. Consistent with the higher stop codon readthrough observed in the lung compared to the brain, the lung exhibited more stop codon readthrough products than the brain, relative to the accurate products, which were used as a loading control (Figs. 2C, and S3A). Furthermore, we compared the mRNA ratio of Halo vs SNAP and Fluc vs Rluc to examine if the low Fluc activity is due to less Fluc RNA region. Both the ratio of Halo/SNAP and Fluc/Rluc RNA regions showed similar levels between organs (Fig. S3B). These results support that the observed differences in stop codon readthrough products in our knock-in mice are due to differences in translation fidelity between organs. Our findings using this mouse model align well with previous studies showing vulnerability to increasing translation errors in the brain and muscles ^27,28^. Thus, reducing mRNA translation errors in the brain and muscles might have a crucial role in ensuring functional proteomes in these organs.

### The brain has low stop codon readthrough genome-wide

To further extend our results from our reporter construct to endogenous mRNA transcripts, we examined the prevalence of stop codon readthrough errors genome-wide using Ribo-seq ^53^, which monitors translation efficiency by sequencing ribosomal footprints at the codon resolution on actively translated transcripts ^9,54^. We leveraged an existing Ribo-seq data set from a previous study that monitored mRNA translation in various organs of 7-week-old young adult mice ^55^. We re-examined this data, evaluating ribosome occupancy and periodicity immediately after the stop codon in the 3’ UTRs of transcripts to estimate the rate of readthrough errors genome-wide. Consistent with our knock-in mouse reporter results, the Ribo-seq data revealed that the brain has fewer ribosomal footprints immediately after the UGA stop codon when compared to the heart, kidney, liver, and lung, indicating less stop codon readthrough from endogenous transcripts (Fig. 2D). Furthermore, codon triplet periodicity was observed in the 3’ UTRs from the kidney and liver and weakly in heart and lung samples, providing additional evidence that these reads resulted from stop codon readthrough and 3’ UTR translation. We also observed similar trends in the UAG and UAA stop codon transcripts (Fig. S3C). Altogether, our knock-in reporter mice and Ribo-seq analyses demonstrate that the brain has the lowest stop codon readthrough rate in 2-month-old mice.

### Stop codon readthrough errors are reduced across development

While young adults exhibited different stop codon readthrough rates in each organ, the E9.5 embryo exhibited an error rate similar to mESCs in all regions. Hence, we reasoned that translation errors would decrease in each organ between E9.5 and young adult mice, particularly across embryonic development, a time in which cells are differentiating, and tissues are patterning to produce functional, complex organs. To further investigate when the stop codon readthrough rate is reduced during embryonic development, we microdissected individual organs from embryos at later developmental stages, such as E11.5, E13.5, and E18.5. We observed a gradual decline in stop codon readthrough rates in the brain and heart across time points when compared to mESCs (Figs. 3A-B). Some tissues, such as the liver and lung, had more consistent stop codon readthrough across embryonic development (Figs. 3C-D). These data suggest that organ-specific translation error rates are gradually established across embryonic development, rather than by a quick switch from low to high fidelity during maturation. We also noticed that translation errors are not exclusively reduced during embryonic development. In the brain and heart, there are differences in stop codon readthrough rate between the day before birth, E18.5, and the mature young adult stages (2-months). Most strikingly, stop codon readthrough in the skeletal muscle at E18.5, specifically quadriceps, as well as developing limb bud is closer to that of mESCs, but stop codon readthrough error in quadriceps is reduced to a similar level to the brain at 2 months (Fig. 3E). This data indicates that organ-specific translation error rates can be established at different time scales depending on the organ. Thus, our research provides the first look into the spatiotemporal landscape of translation fidelity across embryonic development. Although errors during mRNA translation represent a source of stochastic noise in gene expression, our results indicate that the amount of noise can be developmentally programmed, which might have an underappreciated function for cell differentiation and organ patterning across development. Especially, the brain needs to express exceptionally long proteins such as piccolo and huntingtin, which underpin synaptic architecture and axonal function. These findings suggest translation fidelity as a previously unrecognized layer of gene regulation that ensures the production of an organ-specific proteome with such long proteins.

**Fig. 3:**
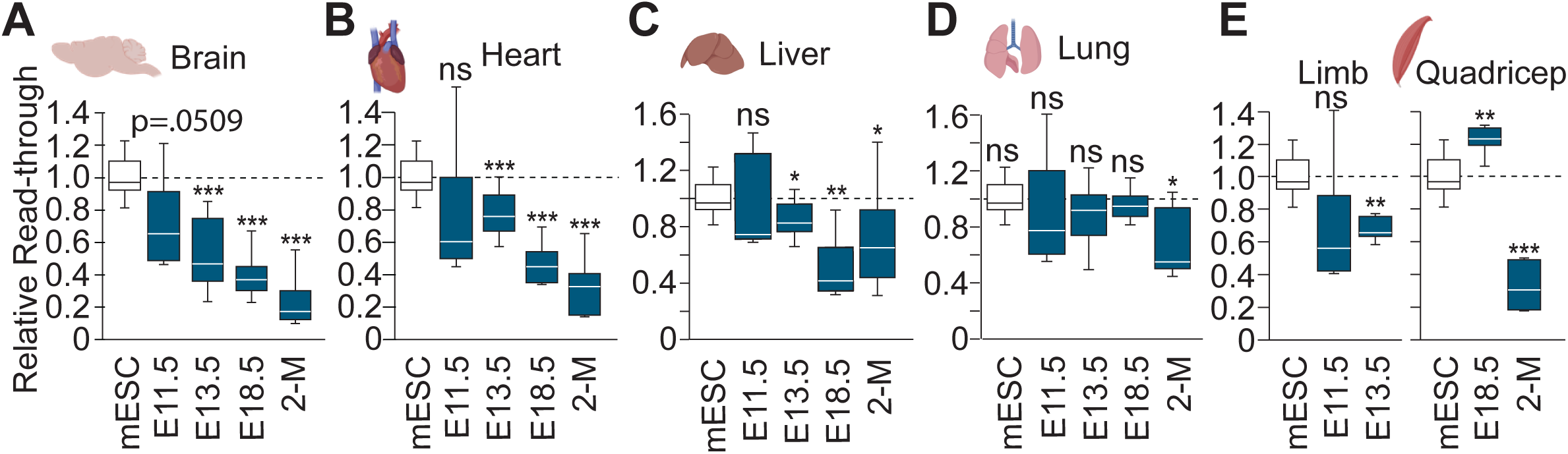
Organ-specific error rates are programmed during embryonic development. (**A** to **E**) Each organ exhibited a different timeframe to develop an organ-specific stop codon readthrough rate. The relative amount of stop codon readthrough to mESCs was quantified throughout each organ’s development in E11.5, E13.5, E18.5, and 2-month-old (2-M). n≥6 harvested from more than 2 litters. Box plots show the interquartile range (IQR) with the median indicated by a central line; whiskers extend to 1.5 × IQR. ns: not significant, *P<0.05, **P<0.01, ***P<0.001 by Mann-Whitney U test.

### Translation errors impair cerebral organoid formation

Based on our observation that stop codon readthrough is gradually reduced during brain development, we next examined the impact of increasing translation errors during brain development using the cerebral organoid model, which reproduces the three-dimensional organization, cell-type diversity, and developmental signaling environment of the brain (Fig. S4A) ^56,57^. Throughout cerebral organoid formation, translation errors were pharmacologically induced by the aminoglycoside paromomycin ^57,58^. Indeed, paromomycin treated organoids increased stop codon readthrough (Fig. 4A). Treated organoids exhibited a smaller size than untreated ones after neuroectoderm induction on day 5. This size difference became more visually prominent by day 9, following the neuron differentiation step, and was maintained thereafter (Figs. 4B, and S4B). This organoid size reduction is consistent with the microcephaly phenotype observed in patients with mutations that reduce translation fidelity ^59–62^.

**Fig. 4:**
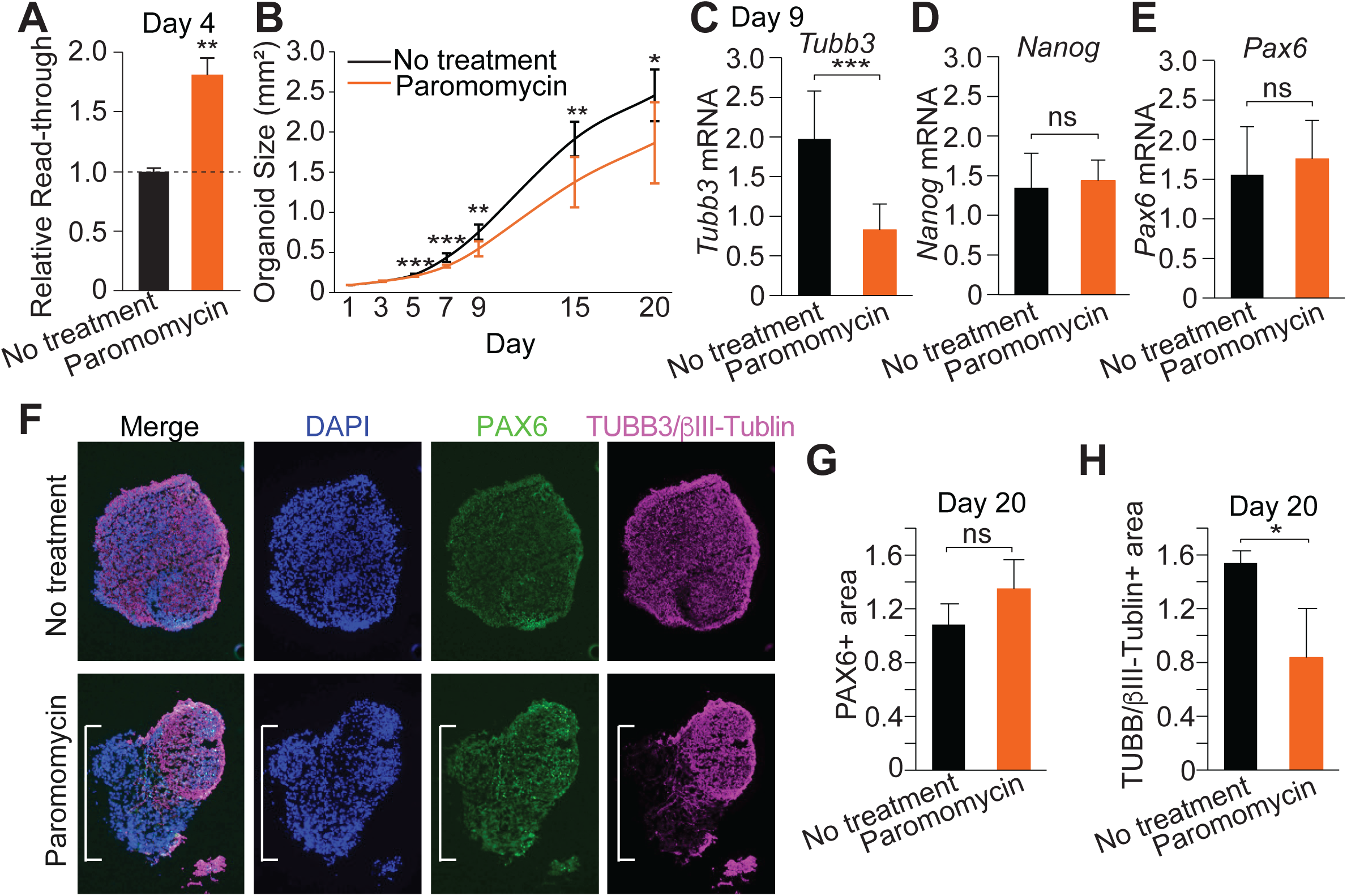
Induced translation errors disrupt cerebral organoid formation. (**A**) Paromomycin treatment increases stop codon readthrough in cerebral organoids on day 4. (**B**) Paromomycin-treated cerebral organoids exhibited a smaller size. (**C** to **E**) Paromomycin-treated day 9 cerebral organoid reduces expression of the mature neuron marker, *Tubb3* mRNA (C), but not the stem cell marker *Nanog* mRNA (D) and neural progenitor cell (NPC) marker *Pax6* mRNA (E). Each mRNA was normalized by *Actb* mRNA. (**F**) Paromomycin treatment reduced the mature neuron population in the cerebral organoid at day 20. Distribution of NPCs and mature neurons with paromomycin and no treatment was visualized by immunofluorescence against PAX6 and TUBB3/βIII-tubulin proteins in frozen section. (**G** to **H**) Paromomycin treatment reduced the neuron marker positive area (H) but did not reduce the NPC marker positive area (G). Each area was normalized by the DAPI area. Bars represent mean ± s.d. of n≥3. ns: not significant, *P<0.05, **P<0.01, ***P<0.001 by Mann-Whitney U test.

To examine if increasing translation errors affects cell population and differentiation, we performed RT-qPCR on day 9 organoids. Interestingly, *Tubb3* mRNA, which is a marker for neuronal differentiation, was reduced under paromomycin treatment using *Actb* mRNA as a normalizer (Fig. 4C). In contrast, the expression of key transcription factors for pluripotent stem cells, *Nanog* mRNA, and neural progenitor cells (NPCs), *Pax6* mRNA, were not affected by paromomycin treatment (Figs. 4D-E). To further assess the consequences of paromomycin treatment and interrogate the effect of the observed mRNA expression changes on the cell population, we performed immunofluorescence staining to examine the abundance and spatial distribution of neuronal cells on day 20. Consistent with our qPCR results from day 9, the distribution and area size of PAX6 positive cells were consistent between paromomycin treated and untreated organoids, indicating the population of NPCs was not affected by increasing translation errors (Figs. 4F-G). Interestingly, paromomycin treated organoids exhibited greater TUBB3/βIII-tubulin negative areas, while untreated organoids indicated equal distribution across the section area. Thus, both the distribution and area size of TUBB3/βIII-tubulin positive neuron cells were markedly reduced under paromomycin treatment (Figs. 4F-H). Given that the PAX6-positive progenitors remained unchanged, our results indicate that increasing translation errors induces a loss of differentiation from NPCs to neuronal cells. Although this phenotype could reflect expansion of radial glia, the reduced organoid size and unchanged PAX6-positive progenitor area argue against this possibility, because radial glial expansion is typically associated with increased progenitor proliferation or delayed neurogenesis ^63^. Instead, these data support impaired neuronal differentiation as the primary consequence of paromomycin treatment. Altogether, these results suggest that increasing translation fidelity during brain development has an important role in neuronal differentiation.

### High translation fidelity facilitates neuron maturation

Our knock-in mice revealed that the brain possesses the lowest stop codon readthrough among the organs tested. Since increased translation errors affect neuron differentiation, we hypothesized that neurons not only have less stop codon readthrough but higher translation fidelity more generally. To examine this hypothesis, we sought to monitor different types of translation errors in neurons. However, monitoring translation errors in the neuron is challenging because of the low efficiency of transient transfection. To overcome this challenge, we generated mESC lines integrating our fidelity reporters into the genome via the PiggyBac transposase system (Fig. S2C). Using these mESC lines, embryoid bodies (EBs) were formed and treated with retinoic acid (RA) to reproducibly produce mature neurons, as evidenced by Tuj1 immunostaining for the neuronal marker βIII-tubulin (TUBB3) (Fig. 5A) ^64^. Because RA-induced neuronal differentiation from EBs generates a mixed population containing both post-mitotic neurons and less-differentiated, non-neuronal/progenitor cells, this culture provides an opportunity to compare translation fidelity between neuron-enriched and non-neuronal, progenitor-enriched fractions within the same genetic and environmental background. These two cell populations were fractionated based on their differences in cell adhesion properties (Fig. 5B). We monitored stop codon readthrough rate using the same construct as our knock-in mouse and found that the non-neuron cell fraction exhibited a higher error rate than that of the neuron fraction (Fig. 5C). This result is consistent with the low stop codon readthrough observed in the brain of our knock-in mice (Fig. 2).

**Fig. 5:**
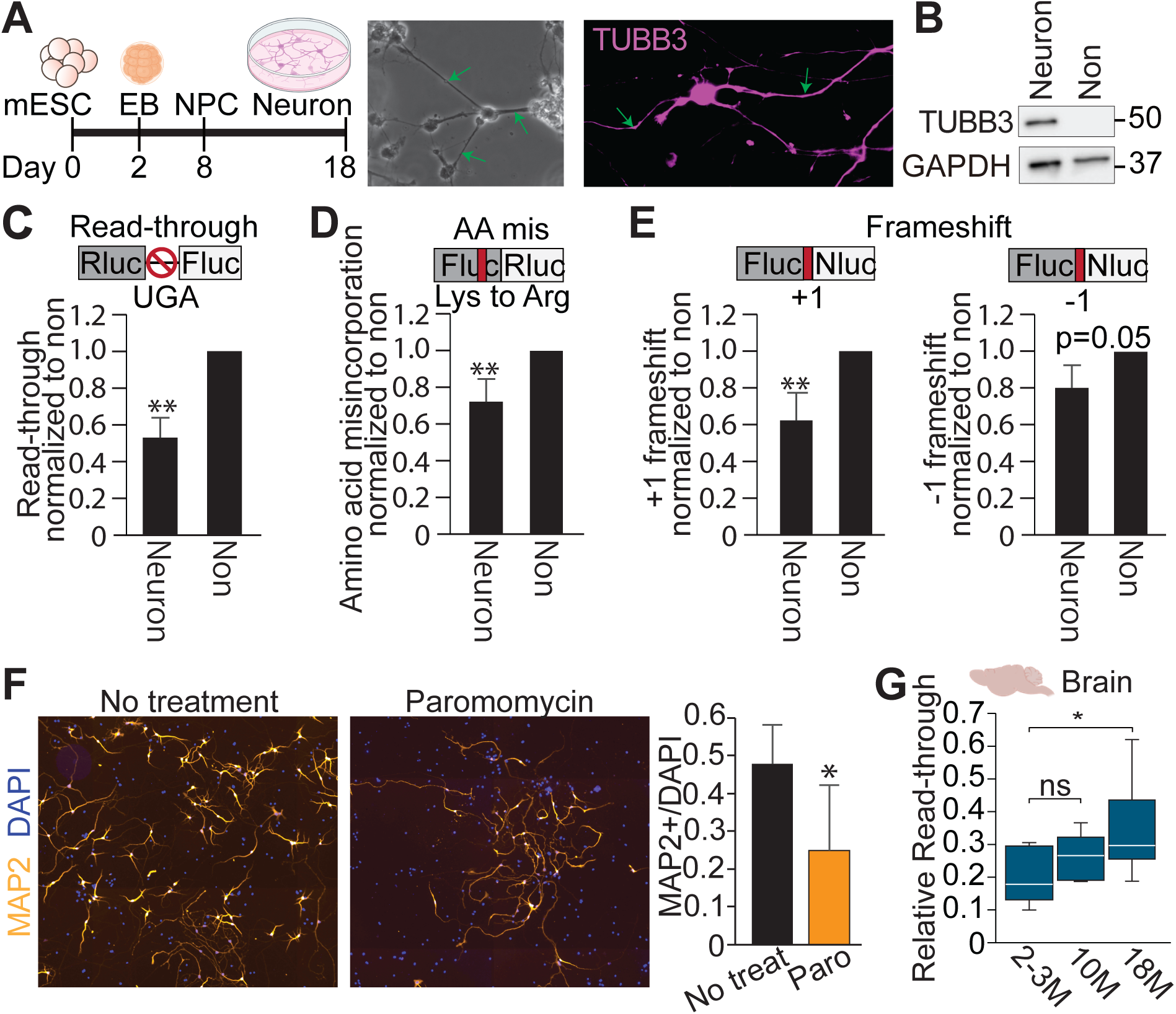
High translation fidelity facilitates efficient neuron maturation. (**A**) Schematic of in vitro neuron differentiation through embryoid body (EB) and neural progenitor cell (NPC) to neuron. Representative image of differentiated neuron in bright image and immunofluorescence using Tuj1 antibody against neuron marker TUBB3/βIII-tubulin protein. Green arrows indicate axons. (**B**) Neuron and Non-neuron (Non) fractions on day 18 of in vitro differentiation. Western blot against TUBB3/βIII-tubulin and GAPDH proteins indicated clear fractionation. (**C** to **E**) Neurons have lower error rates than non-neuron fractions in stop codon readthrough (C), amino acid misincorporation (D), and ribosomal frameshifting (E). Different types of translation errors were quantified using mESCs lines that have genome integration of different dual-luciferase gain-of-activity reporters. Bars represent mean ± s.d. of n≥3. *P<0.05, **P<0.01 by paired t-test. (**F**) Paromomycin treatment inhibits neuron maturation from NPCs derived from human induced pluripotent stem cells (iPSCs). Mature neuron marker MAP2 stains axons in differentiated neurons. Quantification indicated the cell population of MAP2+ cells in three independent iPSC lines (Fig. S5B). Bars represent mean ± s.d. of n≥3. *P<0.05 by paired t-test. (**G**) Stop codon readthrough errors gradually increased in aged brains. The relative stop codon readthrough rate compared to mouse embryonic stem cells (mESCs) was quantified from ≥3 males and ≥3 females. Box plots show the interquartile range (IQR) with the median indicated by a central line; whiskers extend to 1.5 × IQR. ns: not significant, *P<0.05 by Mann-Whitney U test.

In addition to stop codon readthrough, amino acid misincorporation and ribosomal frameshifting were also monitored. To detect low amounts of error product, we tried to improve our reporter constructs using nano luciferase (Nluc). We found that the Fluc-Nluc construct helps to detect low abundance error products compared to other constructs, including Rluc-Fluc, Nluc-Fluc, or bidirectional promoter-driven Fluc and Nluc cassette (Fig. S5A) ^65^. However, this new reporter system could not be applied to amino acid misincorporation because of the lack of a single amino acid substitution that completely kills the luciferase activity. Thus, a traditional point mutation in the active site of Fluc at Lysine 529 to Arginine was used for amino acid misincorporation, which kills activity ^20,66^. Amino acid misincorporation can be monitored by Fluc activity recovered only when the ribosome misincorporates the original amino acid from Arg (AGA) rare codon, to Lys (AAA). Thus, amino acid misincorporation can be monitored by the ratio of Fluc to Rluc activity. Furthermore, ribosomal frameshift was monitored by Fluc-Nluc reporters that contains a +1/-1 frameshift element between Fluc and Nluc. Thus, these reporters will produce active Nluc only when the ribosome makes a frameshift in either the +1 or −1 direction. Combining these reporter systems with in vitro neuron differentiation, neuron cells exhibited less errors than non-neuron cells for amino acid misincorporation and +1 frameshifting (Figs. 5D-E). Although −1 frameshift was not significant, it also indicated a trend in the same direction (p=0.0501) (Fig. 5E). These results demonstrate that neurons have greater translation fidelity in general.

To further gain insight into whether increasing translation errors directly impacts neuron differentiation, we took advantage of a sophisticated induced Pluripotent Stem Cell (iPSC) system, which was already differentiated into NPCs. When we induced translation errors during the maturation step of in vitro neuron differentiation, we observed fewer MAP2 positive mature neurons in iPSC lines produced from different individuals (Figs. 5F, and S5B). This result indicates that the role of translation fidelity in neuron development is conserved in humans, which is consistent with phenotypes observed in patients with translation fidelity defects ^59–62^. Given the sensitive nature of neurons to the proteotoxic stresses during aging and neurodegeneration, establishing higher translation fidelity during neuron and brain development might be crucial to reduce proteostatic stresses across the entire lifespan.

Indeed, increasing UGA stop codon readthrough error in the brain across aging has been observed by using mKate2-Fluc knock-in reporter mice, which is similar to our mouse. Consistent with this result, our knock-in mice also exhibited a gradual increase in stop codon readthrough from 2 to 10 to 18 months old in the brain (Fig. 5G). Since the basal translation error rate in the brain is extremely low, the brain still maintains a relatively low error rate compared to other organs, even in the aged mice (Fig. S5C). However, neuron morphology presents unique challenges with localized protein synthesis, and such slight increases in translation errors might significantly impact the functionality of the proteome. Indeed, the brain increases disparity between mRNA and protein levels across aging ^67^. This aging-associated fidelity reduction might contribute to such age-dependent decline of protein homeostasis. Altogether, our findings uncovered an underappreciated dynamic program of mRNA translation fidelity across development and revealed the crucial function of this fidelity program for neuronal differentiation.

## Discussion

Although the fidelity of mRNA translation is a crucial aspect of gene expression, the spatiotemporal dynamics of translation fidelity have largely been unexplored, especially at the organismal level. By generating a highly quantitative dual-luciferase reporter knock-in mouse model, we charted the first spatiotemporal map of translation fidelity in developing organisms. Our work indicates that translation error rates are substantially lower in vivo than in cultured cells. Given the more stringent physiological constraints present in organisms compared with standard cell culture conditions, this observation is consistent with the traditional dogma that translation errors impose a burden on cellular homeostasis. At the same time, our analyses uncovered pronounced organ-specific differences in translation fidelity in young adult mice that are progressively established during embryonic development. These findings demonstrate that mRNA translation fidelity is not a rigid biochemical constant but rather a dynamic property of gene expression that is tuned by cellular environment and developmental context. Although we cannot formally exclude minor contributions from rare transcriptional errors, multiple lines of evidence strongly support a translational origin of the observed signals. In particular, independent Ribo-seq analyses provided results consistent with our in vivo reporter analysis, revealing reduced endogenous stop codon readthrough in the brain. Together, our results identify translation fidelity as an underappreciated dimension of gene regulation that may contribute to tissue-specific proteome formation by ensuring accurate synthesis of extremely long and structurally demanding proteins, such as titin in muscle and piccolo in neurons.

Our results further show that differentiated neurons exhibit not only lower stop codon readthrough but also reduced amino acid misincorporation and ribosomal frameshifting. Consistent with this, induced translation errors in cerebral organoids and differentiating human iPSCs disrupted neuronal maturation. Although relatively rare, mutations in ribosomal proteins or translation factors that are predicted to reduce translation fidelity have been reported in neurodevelopmental disorders, including microcephaly and autism ^1,2^. Such ribosomal mutations are categorized into a group of diseases collectively referred to as ribosomopathies, which classically present with severe anemia and reduced protein synthesis ^68^. However, ribosomopathies that affect translation fidelity specifically present with neurological phenotypes, which are distinct from those arising from canonical defects in translation capacity. Increasing translation errors during neuronal maturation might disrupt the balance of protein synthesis and degradation, inducing proteotoxic stress and selectively compromising differentiating neurons. In contrast, high levels of proteasome activity in stem cells ^69^ might help buffer elevated translation error products in mESCs and early embryos. Altogether, these findings highlight the crucial function of a cell-type-specific translation fidelity program for neuronal differentiation and brain formation.

The maintenance of proteostasis requires a tight balance between protein synthesis and degradation, a challenge that is particularly acute in long-lived, postmitotic cells such as neurons and muscle fibers. In this context, our observation that the brain and muscles exhibit high translation fidelity early in life may serve as a protective mechanism to limit proteotoxic stress as these tissues establish and maintain their specialized proteomes. Notably, we and others have observed a gradual increase in translation errors across aging in the brain ^34^. In tandem with decreasing protein degradation activity, this age-dependent increase in error might synergistically contribute to the age-dependent reduction of protein homeostasis and seed non-genetic protein aggregation in disease, such as TDP-43 in ALS and Tau and Amyloid beta in Alzheimer’s disease ^70,71^. Our research delineating the dynamics of translation fidelity provides a baseline of translation errors in normal conditions and represents the foundation to further explore the dynamics of mRNA translation fidelity in health and disease.

## Methods

### Generation of knock-in reporter mice

All mice used in this study were housed at the University of Florida. All animal work was reviewed and approved by the University of Florida Animal Care Services (ACS). University of Florida ACS is accredited by the American Association for Accreditation of Laboratory Animal Care (AAALAC). Mice were housed under a 12-hour light/dark cycle with continuous access to food and water. Stop codon readthrough reporter knock-in mice (C57BL/6J-H11^UGA^) were produced in the C57BL/6 H11 background and generated by integrase-mediated transgenesis at the Transgenic, Knockout, and Tumor Model Center (TKTC) at Stanford University. Age-matched, litter-matched cohorts of transgenic and wild-type were regularly weighed in grams from birth to 22-weeks.

### Cell lines

Low-passage E14 mouse embryonic stem cells (mESCs) were grown in a growth medium composed of knockout DMEM (Invitrogen, 10829-018) with 15% ESC grade FBS (EMD Millipore, ES-009-B), 1x L-glutamine (EMD Millipore, TMS-002-C), 1% Penicillin-Streptomycin (Gibco 15-140-122), 1x non-essential amino acid (EMD Millipore, TMS-001-C), 0.1% 2-mercaptoethanol (Gibco 21985-023), 0.01% mLIF 10^7^ (Gemini Bioproducts 300-365P-100) in 37 C, 5% CO_2_ incubator. C3H/10T/1/2 and NIH3T3 cells were grown under standard conditions in DMEM (Invitrogen, 11965-092), 10% FBS, and 1% Penicillin-Streptomycin. HAP1 cells were grown under standard conditions in IMDM (Invitrogen, 12440-046) supplemented with 10% FBS, and 1% Penicillin-Streptomycin.

Lipofectamine 2000 (Invitrogen 11668019) was used for plasmid transfection. SNAP and Halo tags were visualized by adding 0.2µM JF646 HaloTag ligand (Promega GA1120) and/or 1µM Oregon Green SNAP ligand (NEB S9104) in mESC medium for live imaging.

mESC lines harboring translation fidelity reporters were generated using PiggyBac genome integration. For each cell line, mESCs were transfected with 1µg of reporter plasmid containing PiggyBac recognition sites and 1µg of plasmid for the PiggyBac transposase expression vector, using Lipofectamine 2000 (Invitrogen 11668019). Transfections were performed in a 6-well plate coated with 0.1% gelatin at 1.0 x 10^6^ cells per well in 2 mL of growth media. After 24 hours of incubation, approximately 1,000 single cells were transferred to a 10cm dish coated with 0.1% gelatin and allowed to grow as single colonies for five to seven days. Once colonies became visible, each colony was transferred to a single well of a 96-well plate and dissociated by trypsin, then further cultured for genotyping and confirmation of luciferase activity before downstream use. We generated at least 3 colonies for each construct.

MEF lines were generated from E13.5 embryos of C57BL/6J-H11^UGA/+^, following standard procedures. Briefly, pregnant females were euthanized, and embryos were dissected. Heads and visceral organs were removed, and the remaining carcasses were minced and dissociated by trypsinization (0.05% trypsin-EDTA, 10 min at 37 °C) with periodic pipetting. The cell suspension was filtered through a 70-µm strainer and plated onto dishes in MEF medium (DMEM supplemented with 2 mM L-glutamine, 10% FBS, and penicillin–streptomycin). MEFs were expanded for one or two passages and cryopreserved for subsequent experiments. Cells were seeded on the 24-well plate with or without 24hr treatment of 1mg/ml paromomycin to monitor stop codon readthrough rate.

### Tissue harvest from mice and mouse embryos

Tissue samples were harvested from both adult mice and embryos. Adult mice were anesthetized via isoflurane induction, then intracardiac perfusion was performed with 1X PBS to remove blood contamination from organs. Following perfusion, the brain, quadriceps, heart, lung, kidney, and liver were immediately extracted and flash frozen with liquid nitrogen and stored at −80 °C until lysis. Mouse embryos were collected at E9.5, E11.5, E13.5, and E18.5 from pregnant dams. Pregnant mice were sacrificed by CO_2_ euthanasia followed by cervical dislocation. The uterus was then dissected out and placed in 1X PBS for microdissection of the embryonic organs. Organs were flash frozen with liquid nitrogen and stored at −80 °C until lysis.

### Luciferase assay

Frozen organ samples from adult mice were ground with pre-chilled mortar and pestle under liquid nitrogen. Powderized lysate was resuspended in 1x Passive Lysis Buffer (Promega, Cat PR-E1941a) and centrifuged before assay. Embryonic organs were resuspended in 1x Passive Lysis Buffer then ground with pestle in 1.7mL tubes and centrifuged before assay. For cell culture samples, media was aspirated from cell cultures followed by a 1x PBS wash, then incubated in 1x Passive Lysis Buffer for >5 minutes at room temperature. Cell lysates were then collected and stored at −80 °C until assay. In vitro differentiated neurons were separated from non-neuronal cells prior to cell lysis via differential timing of 1x Trypsin-EDTA (Life Technologies, 15400-054) whereby 2 minutes of incubation for non-neuronal and 3 more minutes for neurons. Trypsin was inactivated by adding media. Cells were centrifuged and resuspended in 1x Passive Lysis Buffer, then stored at −80 °C until assay.

Prior to performing the luciferase assay, lysates were centrifuged at 1000rpm for 3 minutes at 4 °C. Afterwards, 10uL of each lysate was loaded into a well of black luciferase assay plates (Fisher Scientific, TPB9096) and assayed using the Dual-Luciferase® Reporter Assay System (Promega, Cat E1910) or Nano-Glo® Dual-Luciferase® Reporter Assay System (Promega, Cat N1610). To measure mistranslation, tandem reporters of Rluc and Fluc or Fluc and Nluc were used by which the first luciferase’s activity in the reporter was used to normalize the activity of the second after accounting for background luminescence.

### RT-qPCR

RNA from organoids was extracted using TRIzol (Invitrogen, Cat. 15596026). RNA from organs was extracted using the Quick-RNA MiniPrep kit (Zymo, 11-327) with Turbo DNase treatment (Ambion, AM2238) supplemented with SUPERaseIn (Invitrogen, AM2694), followed by column purification with the RNA Clean & Concentrator −5 kit (Zymo, R1016). 100ng of RNA was reverse transcribed using iScript Reverse Transcription Supermix (Bio-Rad, Cat. 1708841), and qPCR was performed with primers listed in Table S2 and SsoAdvanced Universal SYBR Green Supermix (Bio-Rad, Cat. 1725274) using the CFX Opus 384 Real Time PCR System (BioRad, 12011452). RNA levels were quantified using a standard curve method by CFX manager (Bio-Rad).

### Immunoprecipitation and western blotting

Whole brain and lung tissues from transgenic and wild-type mice were flash-frozen in liquid nitrogen and pulverized using a pre-chilled mortar and pestle under liquid nitrogen. The resulting tissue powder was resuspended in immunoprecipitation (IP) buffer (50mM Tris-HCl, pH 7.5; 150 mM NaCl; 2.5mM MgCl_2_; 1% Triton X-100) supplemented with 20U/mL TURBO DNase (Invitrogen, AM2238), 200U/mL RNaseOUT Ribonuclease Inhibitor (Invitrogen, 10777019), and 1x Halt Protease and Phosphatase Inhibitor Cocktail (Thermo Scientific, 78442), using 1mL for brain samples and 3mL for lung samples. Lysates were incubated on a rotating platform at 4°C for 30 min. Debris was removed by centrifugation at 1,800 × g for 5 min at 4°C, followed by a second centrifugation of the supernatant at 15,000 × g for 5 min at 4°C. The resulting supernatant was retained as input for Western blot analysis. Immunoprecipitation was performed using ANTI-FLAG M2 Affinity Gel (Millipore, A2220) pre-equilibrated with IP buffer. Lysates were incubated with 20μL net volume of FLAG beads for 1.5 h at 4°C with gentle rotation. Following incubation, the bead–lysate slurry was transferred to columns. Beads were washed with 3mL of IP buffer, followed by 2mL of 1x PBS. For elution, beads were incubated at 4°C for 10 min in 20μL of 1x PBS containing 100μg/mL FLAG peptide for FLAG IP, followed by centrifugation at 13,000 × g for 1 min at 4°C to collect the eluate. A second elution was performed using the same procedure. The two eluates were combined.

For Western blotting, tissue samples from FLAG IP were loaded onto 4–20% Mini-PROTEAN TGX Precast Protein Gels (Bio-Rad, 4561094) based on equal Rluc activity. Proteins were transferred to PVDF membranes using the Trans-Blot Turbo system (Bio-Rad, 170-4273) according to the manufacturer’s instructions. Reporter proteins were visualized by anti-FLAG antibody (Proteintech, 20543-1-AP). Signal detection was performed using Clarity Max Western ECL Substrate (Bio-Rad, 1705062S) and imaged with the ChemiDoc MP Imaging System (Bio-Rad, 17001402). Western blotting for Tuj1 in neuron and non-neuron lysates was performed in the same way, except that the Tuj1 antibody was used (Millipore Sigma, T2200).

### Ribosome profiling data analysis

Ribosome profiling data for mouse brain and other tissues were obtained from a publicly available dataset from 7-week-old mice ^55^. Raw sequencing data were processed using the RiboFlow pipeline which performs adapter trimming, contaminant filtering, alignment to the mouse GRCm39 reference genome, and read length–specific quality control ^72^. Following preprocessing, A-site offsets were calculated independently for each sample and each read length (28–32 nt). These offsets were applied to align reads with the ribosomal A-site. Metagene periodicity profiles around annotated stop codons were produced.

### Cerebral organoid formation

Cerebral organoids were generated as described elsewhere ^73^. In brief, 2×10^3^ mESCs were seeded into each well of a 96-well U-bottom bottom plate in 100µL of seeding medium, defined as 10% KnockOut Serum Replacement (Gibco, 10828010), 2 mM L-glutamine, 1 mM sodium pyruvate (Gibco, 11360070), 0.1mM ES Cell Qualified MEM Non Essential Amino Acids, 0.1mM 2-ME, and 10μM SB431542 in G-MEM (Gibco, 11710035). Following four days of embryoid body formation, with media change every other day, media was exchanged with neuroectoderm induction medium, defined as DMEM/F-12 supplemented with 1% N-2, 1% GlutaMax, 1% MEM-NEAA, and 1ug/ml Heparin (Sigma, H3149-100KU). After two days of neuroectoderm induction, embryoid bodies were embedded in droplets of Matrigel Matrix for Organoid Culture (Corning, 08-774-406) and grown in differentiation medium, defined as a 1:1 mixture of DMEM/F-12 and Neurobasal medium with 1% B-27 minus vitamin A (Gibco, 12587010), 1% GlutaMax (ThermoFisher, 35050061), 0.5% N-2, 3.5 µl/L 2-ME, and 1:4000 human insulin (Sigma, 11061680). After five days of growth in differentiation medium, with media change every other day, media was exchanged with differentiation medium made from B-27 supplement, rather than B-27 minus vitamin A. Organoids were placed on a shaker at 65 rpm, and the media was changed every two to three days until harvest. For paromomycin supplementation, 1mg/mL paromomycin was supplemented in organoid media during each media change. Surface area measurements were taken every other day for 9 days, then at days 15 and 20 via bright field imaging with the EVOS M5000 microscope (Invitrogen, AMF5000). For surface area measurements, the microscope’s native measuring tools were used to measure outlines of each organoid.

### In vitro neuron differentiation from mESC

In vitro differentiated neurons were generated as described elsewhere ^64^. In brief, 2×10^3^ mESCs were seeded into each well of a 96-well U-bottom plate in 100µL of basal differentiation medium, defined as 15% ES Cell Qualified FBS (Millipore Sigma, ES009-M), 1% Penicillin-Streptomycin (Gibco, 15-140-122), 1% ES Cell Qualified MEM Non Essential Amino Acids (Fisher Scientific, TMS-001-C), and 0.1mM 2-ME (Gibco, 21985-023) in DMEM/F-12 (Gibco, 11320-033). After two days of embryoid body formation, 50 embryoid bodies were transferred to each well of a 6-well plate coated with 0.1% gelatin. At this stage, basal differentiation medium was supplemented with 1µM retinoic acid (Sigma, R2625). After six days of retinoic acid supplementation, embedded embryoid bodies were dissociated with 1X Trypsin-EDTA (Life Technologies, 15400-054), and 1×10^5^ of the resulting cells were seeded in each well of a gelatinized 12-well plate. Six hours after seeding, media was exchanged with a 1:1 mixture of DMEM/F-12 and Neurobasal medium (Invitrogen, 21103-049) supplemented with 2% B-27 (Gibco, 17504044), 1% N-2 (Invitrogen, 17502-048), 1% GlutaMAX (Gibco, 35050061), and 0.1mM 2-ME. Media was changed every other day for 10 days.

### Neuron differentiation from iPSC-derived Neural Progenitor Cells (NPCs)

Three NPC lines derived from iPSC lines were produced from healthy individuals. These NPCs were in vitro differentiated into neurons following the protocol associated with STEMdiff Forebrain Neuron Differentiation Kit (STEMCELL Technology, 08600). iPSCs were differentiated into NPCs by following the STEMdiff Neural Induction kit (STEMCELL Technology, 08581), which uses SMADi inhibition to block TGF-β/BMP-dependent SMAD signaling for efficient neural induction. NPCs were passed several times after thawing in NPC medium (STEMCELL Technology, 05833). Then, NPCs were seeded with high density (0.1×10^6^ cells/cm^2^) and cultured for 6 days under STEMdiff Forebrain Neuron Differentiation medium (STEMCELL Technology, 08600). Differentiated cells were seeded with lower density (1×10^4^ cells/cm) and cultured 20 days for maturation under Brain Phys Neuronal Medium (STEMCELL Technology, 05790) supplemented with 20ng/ml B27 Neuronal Supplement, 1x N2 Supplement-A, 20ng/ml Human Recombinant BDNF, 20ng/ml Human Recombinant GDNF, 1mM Dibutyryl-cAMP, 200nM L-Ascorbic Acid, and Brainphys.

### Immunofluorescence staining

Neurons and cerebral organoids were stained for immunofluorescence to assess neuronal development. Prior to staining, samples were fixed in 4% paraformaldehyde for 20 minutes at 4°C. Additionally, organoids were allowed to sink in 30% sucrose overnight, embedded in Optimal Cutting Temperature Compound (SAKURA FINETEK, 4583), frozen, and sectioned at 10µm. Samples were permeabilized in IF buffer, defined as 1% normal goat serum (MP Biomedicals, 0929391-CF), 1% bovine serum albumin (Sigma, A9647), and 0.5% Triton X-100 (Sigma, T8787-250ML) in 1X PBS. Blocking was done in IF buffer at a Triton X-100 concentration of 0.025%, and antibodies were diluted in this solution as well. Primary antibodies included 1:100 PAX6 (DSHB, AB_528427), 1:1000 Anti-β-Tubulin III Antibody (Sigma, T2200), MAP2 (Invitrogen, PA1-16751). Secondary antibodies included 1:1000 Goat anti-Mouse IgG (H+L) Superclonal Secondary Antibody, Alexa Fluor™ 488 (Invitrogen, A28175) and 1:1000 Goat anti-Rabbit IgG (H+L) Highly Cross-Adsorbed Secondary Antibody, Alexa Fluor 647 (Invitrogen, A21245). Slides were mounted with Fluoromount-G (Southern Biotech, 0100-01) and examined using the EVOS M5000 microscope (Invitrogen, AMF5000) or REVOLUTION microscope (ECHO). At least 3 sections were used for each organoid condition or neuron culture. Quantification of antibody area was performed using QuPath. For each section, the DAPI+ area was used for normalization.

### Statistical analysis

A minimum of 3 biological replicates were used for each group in each experiment. For mouse experiments, at minimum of 3 mice from each sex was used in each group, and the average of technical replicates was used to represent each mouse. The Mann-Whitney U test or Student’s T test was used to test differences between two groups as described in legends. A p-value of less than 0.05 was deemed significant.

## Acknowledgments

We thank members of Fujii lab, Nguyen lab, and the Center for NeuroGenetics for critical discussions and Dr. Olena Zhulyn for proofreading of the manuscript. We thank Stanford Transgenic, Knockout, and Tumor Model Center for generating knock-in mouse. We thank BioRender for organ cartoons used in our Figures.

## Author contributions

Conceptualization: K.F.

Methodology: J.A.B., P.A., I.T., A.S., L.N., and K.F.

Investigation: J.A.B., P.A., I.T., R.K., K.C.M., and K.F.

Bioinformatics: A.B.R.

Visualization: J.A.B., P.A., and K.F.

Supervision: J.A.B., and K.F.

Writing—original draft: J.A.B., I.T., and K.F.

Writing—review & editing: J.A.B., P.A., I.T., A.B.R., R.N., K.C.M., A.S., L.N., and K.F.

## Data and materials availability

All data are available in the main text or the supplementary materials.

