## Supplementary Figures and Tables for "The fidelity of mRNA translation as a novel regulatory layer for brain development"

Jacob A Best et al.

The PDF file includes:

Figs. S1 to S5

Table S1 to S2

### Sup Fig.1

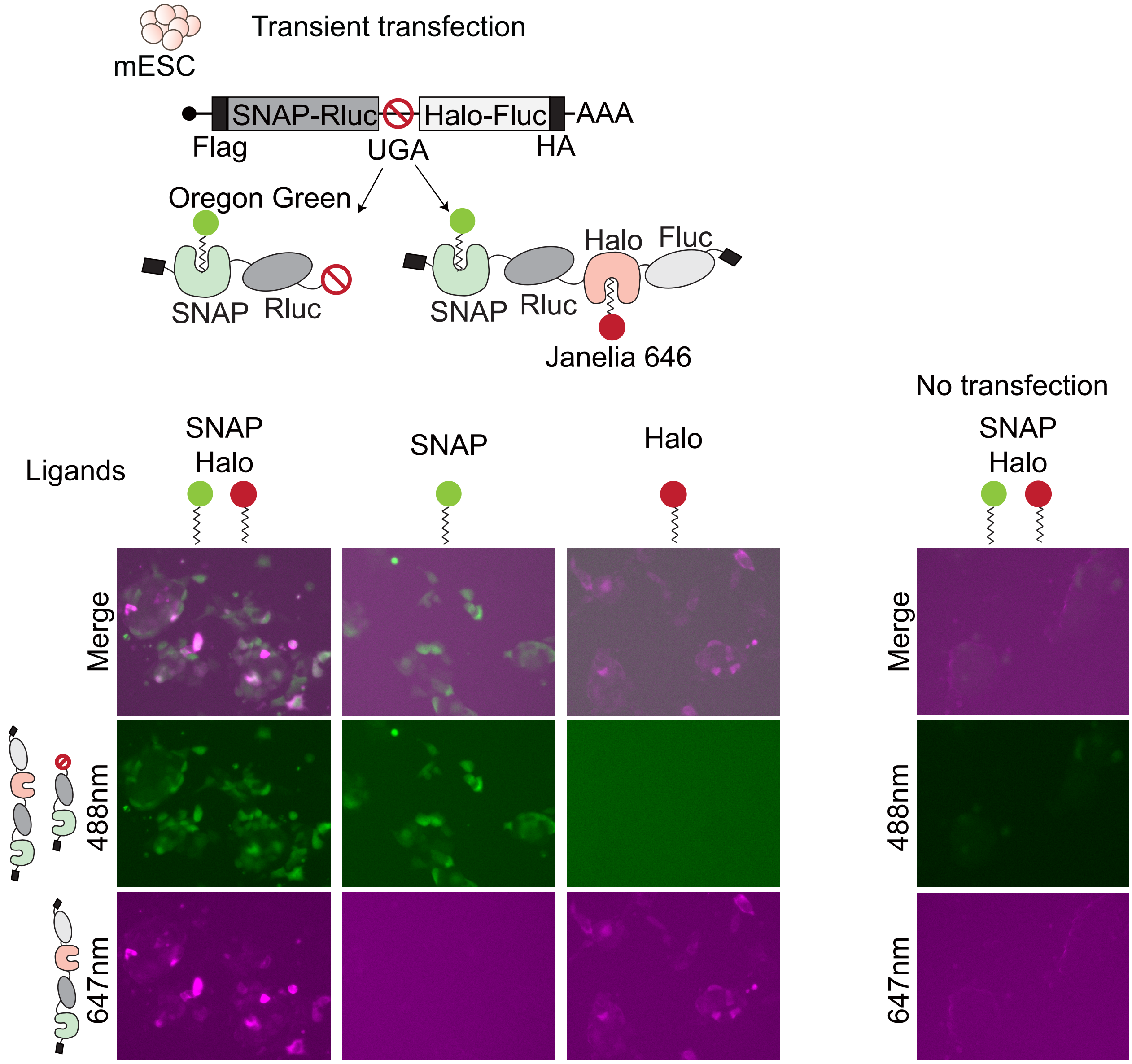

Transient transfection
mESC
SNAP-Rluc
Halo-Fluc
AAA
Flag
UGA
HA
Oregon Green
Halo
Fluc
SNAP
Rluc
SNAP
Rluc
Janelia 646
No transfection
Ligands
SNAP
Halo
SNAP
Halo
SNAP
Halo
Merge
488nm
647nm
Merge
488nm
647nm

**Fig. S1. Visualization of stop codon readthrough derived proteins in transiently transfected live mouse embryonic stem cells (mESCs).** To visualize error products, gain-of-activity reporter with SNAP-Rluc-UGA-Halo-Fluc was transiently transfected into mESCs. All products produced from this construct were visualized by Oregon Green conjugated SNAP ligand. Halo tag containing UGA stop codon readthrough product were visualized by Janelia 646 conjugated ligand. Both SNAP and Halo signals were ligand and product dependent. Each cell indicated surprising heterogeneity for the amount of Read-through products.

### Sup Fig.2

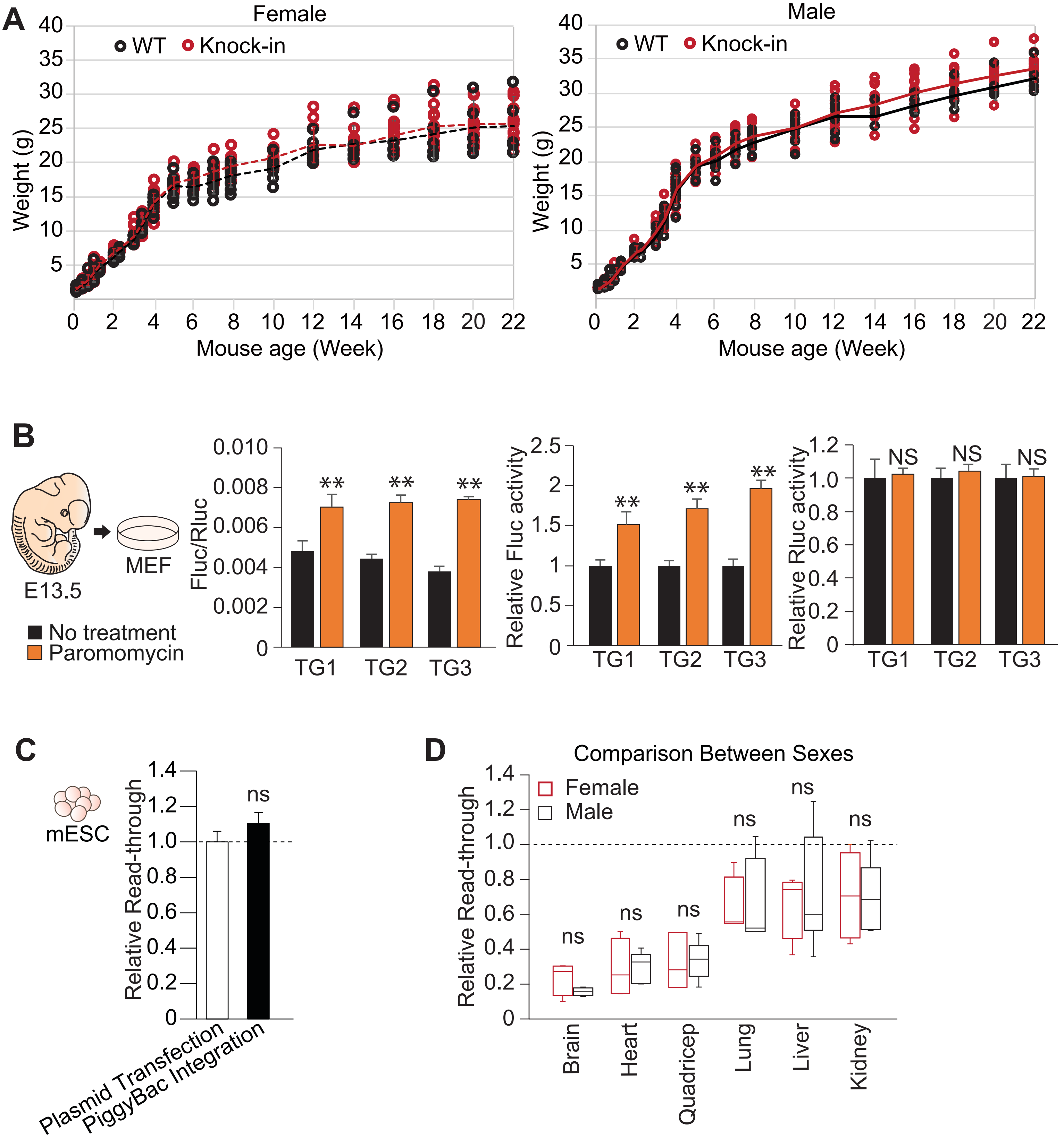

A
Female
WT
Knock-in
Weight (g)
Mouse age (Week)
Male
B
E13.5
MEF
No treatment
Paromomycin
Fluc/Rluc
Relative Fluc activity
Relative Rluc activity
TG1
TG2
TG3
**
NS
C
mESC
Relative Read-through
ns
Plasmid Transfection
PiggyBac Integration
D
Comparison Between Sexes
Female
Male
Brain
Heart
Quadricep
Lung
Liver
Kidney

**Fig. S2. Developing stop codon readthrough reporter knock-in mice.** (**A**) Knock-in mice exhibit growth curves comparable to wild-type littermate controls on a C57BL/6J background. Growth was monitored by weighing from P0 to 22 weeks in both females and males. n>3. (**B**) Challenge of mouse embryonic fibroblasts (MEFs) from E13.5 knock-in mice embryos with paromomycin. Relative stop codon readthrough rate in three independent MEF lines from different litters was monitored with or without paromomycin treatment. Stop codon readthrough rate was calculated as Fluc to Rluc values. Stop codon readthrough products indicated by Fluc activity were increased with paromomycin treatment. Rluc activity did not change with and without paromomycin treatment. Each litter has more than three cell lines. Data are graphed as mean ± SD (N>3). **P<0.01 by unpaired t test. **(C)** The relative readthrough rates in mESCs are comparable between PiggyBac genome integration and transient transfection. Data are graphed as mean ± SD (N>3). NS by unpaired t test. **(D)** The stop codon readthrough rate between females and males did not change in 2-month-old knock-in mice in the brain, heart, quadriceps, lung, liver, and kidney. The relative stop codon readthrough rate in each organ compared to mouse embryonic stem cells (mESCs) was quantified from ≥3 males or ≥3 females. Box plots show the interquartile range (IQR) with the median indicated by a central line; whiskers extend to 1.5 × IQR. ns: not significant by Mann-Whitney U test.

**Sup Fig.3**

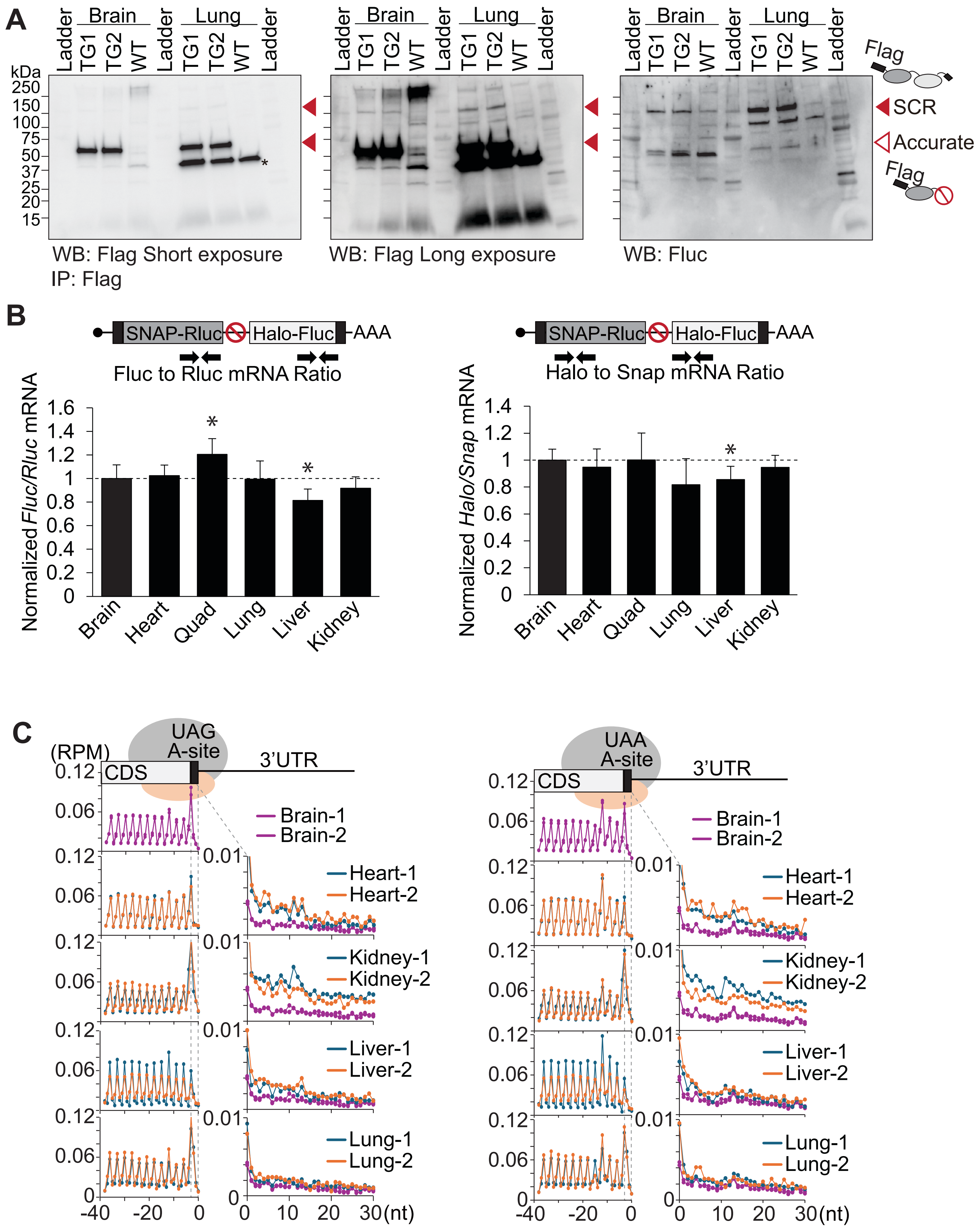
A
Ladder
Brain
TG1
TG2
WT
Lung
kDa
250
150
100
75
50
37
25
20
15
WB: Flag Short exposure
IP: Flag
WB: Flag Long exposure
WB: Fluc
Flag
SCR
Accurate
B
SNAP-Rluc
Halo-Fluc
AAA
Fluc to Rluc mRNA Ratio
Normalized Fluc/Rluc mRNA
Halo to Snap mRNA Ratio
Normalized Halo/Snap mRNA
Brain
Heart
Quad
Lung
Liver
Kidney
C
(RPM)
UAG
A-site
UAA
CDS
3'UTR
Brain-1
Brain-2
Heart-1
Heart-2
Kidney-1
Kidney-2
Liver-1
Liver-2
Lung-1
Lung-2
(nt)

**Fig. S3. The brain has less readthrough products, which was not due to less RNA amount.** (**A**) Reporter derived proteins were purified by Flag immunoprecipitation (IP) and detected by western blotting (WB) using anti-Flag or anti-Fluc. Flag antibody detects both accurate (58 kDa) and stop codon readthrough products (157 kDa). Bands observed in the WT lane are non-specific protein (*). Flag long exposure and Fluc can visualize a low amount of stop codon readthrough products. Consistent with the luciferase assay, lung samples have more readthrough products than the brain sample. (**B**) There is a possibility that the low amount of readthrough products can be due to the fewer RNA molecules containing Halo-Fluc. qPCR analysis comparing RNA between Rluc vs Fluc and SNAP vs Halo indicated that all organs have similar levels of each portion of the transcript as the brain. In contrast to the high error rate observed in the liver, the liver appears to have less error encoding of the transcript portion. Data are graphed as mean ± SD (N=6). (**C**) The brain has a low UAG and UAA stop codon readthrough rate in endogenous transcripts. Ribosome profiling (Ribo-seq) datasets from a previously published 7-week-old young adult mouse show a genome-wide trend of fewer ribosome footprints immediately after the stop codons in the brain compared to other organs. Y-axis plotted in reads per million (RPM).

**Sup Fig.4**

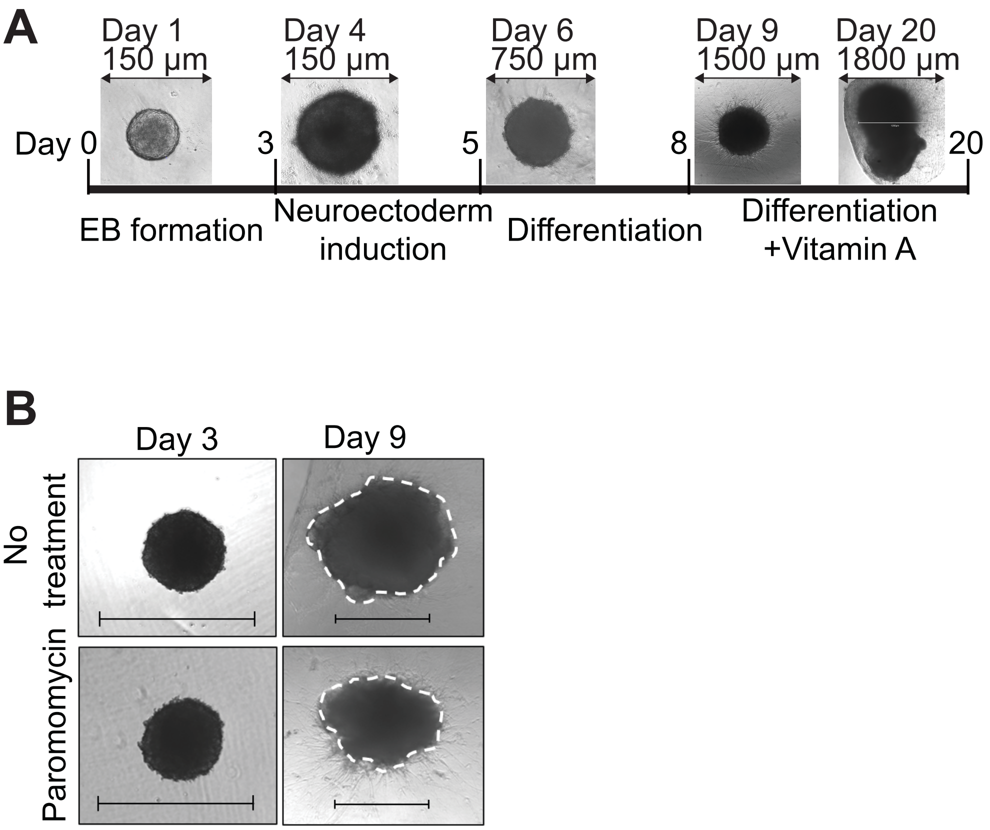
A
Day 1
150 µm
Day 4
150 µm
Day 6
750 µm
Day 9
1500 µm
Day 20
1800 µm
Day 0
3
5
8
20
EB formation
Neuroectoderm induction
Differentiation
Differentiation +Vitamin A
B
Day 3
Day 9
No treatment
Paromomycin

**Fig. S4. Images of cerebral organoids during development.** (**A**) Representative pictures of cerebral organoid generation across days. (**B**) Comparative images of cerebral organoids with and without paromomycin at days 3 and 9 (scale bar=750µm).

**Sup Fig. 5**

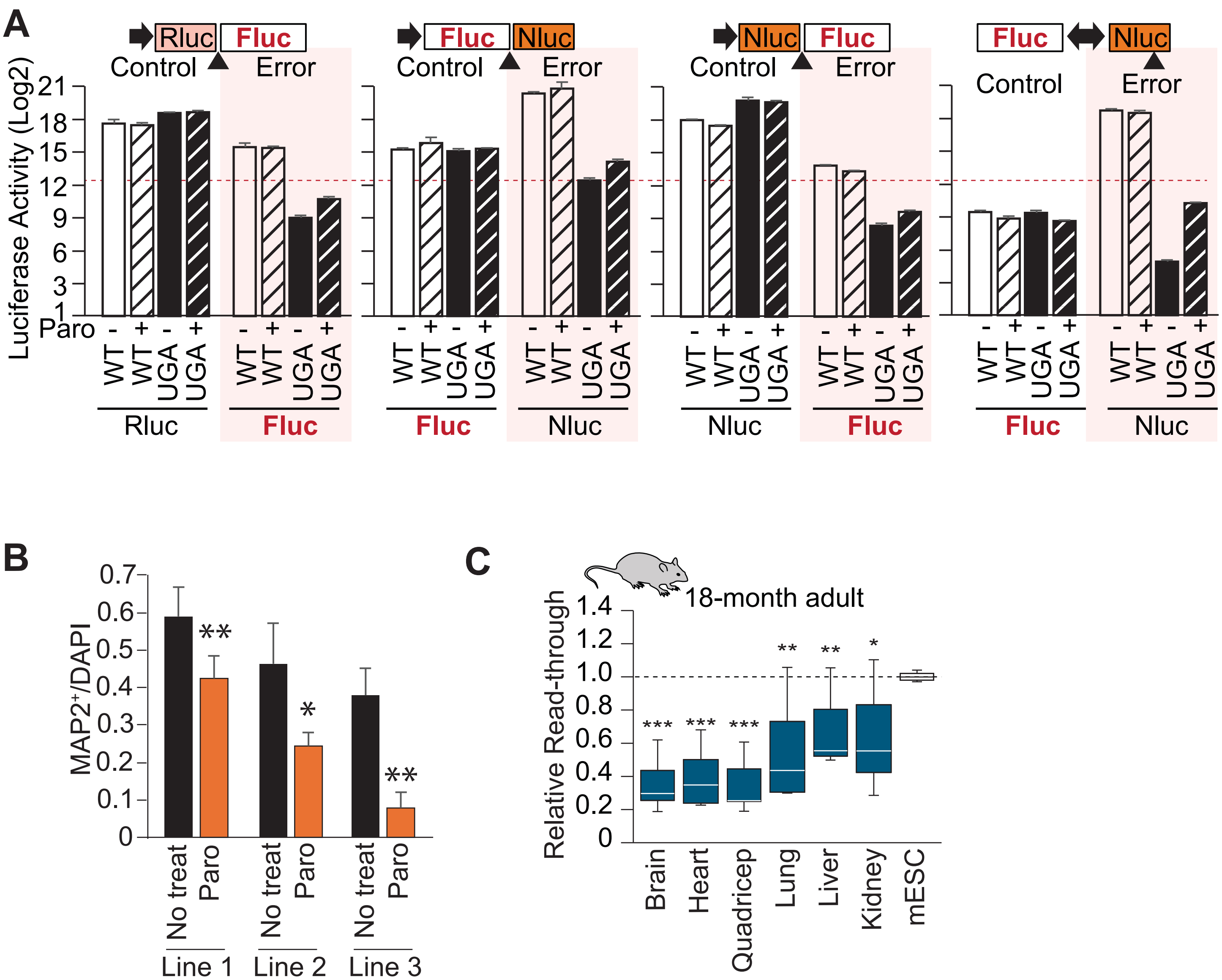

A
Rluc
Fluc
Control
Error
Fluc
Nluc
Control
Error
Nluc
Fluc
Control
Error
Fluc
Nluc
Control
Error
Luciferase Activity (Log2)
21
18
15
12
9
6
3
1
Paro
- + - +
WT
WT
UGA
UGA
Rluc
Fluc
Fluc
Nluc
Nluc
Fluc
Fluc
Nluc
B
MAP2+/DAPI
0.7
0.6
0.5
0.4
0.3
0.2
0.1
0
**
*
**
No treat
Paro
Line 1
Line 2
Line 3
C
18-month adult
Relative Read-through
1.4
1.2
1.0
0.8
0.6
0.4
0.2
0
***
***
***
**
**
*
Brain
Heart
Quadricep
Lung
Liver
Kidney
mESC

**Fig. S5. Establishing reporter-integrated mESCs.** (**A**) Optimizing luciferase combination for the efficient detection of error products. Different combinations of luciferases with UGA stop codon or no-stop (WT) constructs are examined in mESCs. Both under the paromomycin treatment and no treatment have been tested. Detecting errors by Nluc in Fluc-Nluc construct has the highest sensitivity. >3 (**B**) Each individual iPSC line indicated less differentiated MAP2$^{+}$ mature neurons under paromomycin treatment. Data are graphed as mean ± SD (N>3). *P<0.05, **P<0.01 by unpaired t test. (**C**) The relative stop codon readthrough rate in 18-month-old mice in each organ compared to mESCs was quantified from ≥3 males and ≥3 females. In the box plot: Horizontal white lines denote median values; boxes extend from the 25th to the 75th percentile; vertical lines represent the most extreme values within 1.5 interquartile range of the 25th to the 75th percentile of each group. *P<0.05, **P<0.01, ***P<0.001 by Mann-Whitney U test.

Table S1. List of the plasmids used in this study

| Construct | Source | Name |
|---|---|---|
| BT378 CAG Flag-SNAP-Rluc-UGA-Halo-Fluc-HA | This paper | P19 |
| BT378 CAG Flag-SNAP-Rluc-UAG-Halo-Fluc-HA | This paper | P30 |
| BT378 CAG Flag-SNAP-Rluc-UAA-Halo-Fluc-HA | This paper | P31 |
| BT378 CAG Flag-SNAP-Rluc-no stop-Halo-Fluc-HA | This paper | P32 |
| PiggyBac CAG Flag-SNAP-Rluc-UGA-Halo-Fluc-HA | This paper | P40 |
| PiggyBac CAG Flag-Fluc(K223R)-Rluc-HA | This paper | P223 |
| PiggyBac CAG Flag-Fluc-(+1)-Nluc-HA | This paper | P222 |
| PiggyBac CAG Flag-Fluc-(-1)-Nluc-HA | This paper | P221 |
| PiggyBac CAG Flag-Rluc-UGA-Fluc-HA | This paper | P77 |
| PiggyBac CAG Flag-Rluc-no stop-Fluc-HA | This paper | P78 |
| PiggyBac CAG Flag-Fluc-UGA-Nluc-HA | This paper | P81 |
| PiggyBac CAG Flag-Fluc-no stop-Nluc-HA | This paper | P82 |
| PiggyBac CAG Flag-Nluc-UGA-Fluc-HA | This paper | P79 |
| PiggyBac CAG Flag-Nluc-no stop-Fluc-HA | This paper | P80 |
| PiggyBac Flag-Fluc-BiCMV-Nluc(UGA)-HC | This paper | P103 |
| PiggyBac Flag-Fluc-BiCMV-Nluc(WT)-HC | This paper | P104 |

Table S2. List of the qPCR primers

| Primer | Sequence (5' to 3') |
|---|---|
| Nanog-Forward | GAAGCATCGAATTCTGGGAACGC |
| Nanog-Reverse | AGGCAGGTCTTCAGAGGAAGGG |
| Pax6-Forward | CTGAGGAACCAGAGAAGACAGG |
| Pax6-Reverse | CATGGAACCTGATGTGAAGGAGG |
| Tuj1-Forward | CATCAGCGATGAGCACGGCATA |
| Tuj1-Reverse | GGTTCCAAGTCCACCAGAATGG |
| β-Actin-Forward | GCCAACCGTGAAAAGATGAC |
| β-Actin-Reverse | CATCACAATGCCTGTGGTAC |
| Fluc-Forward | CGGGCGCGGTCGGTAAAGT |
| Fluc-Reverse | CCAGCGTTTTCCCGGTATCCA |
| Rluc-Forward | TGGAGAATAACTTCTTCGTGGA |
| Rluc-Reverse | TTGGACGACGAACTTCACC |
| Halo-Fwd | GCGCGTCAAAGGTATTGCAT |
| Halo-Rev | GGGCAAATTCTGGCCATTCG |
| Snap-Fwd | CCTGGCTCAACGCCTACTTT |
| Snap-Rev | AAGCTCTCCTGCTGGAACAC |
